## Supplementary File 1 for "Neuroelectrophysiology-Compatible Electrolytic Lesioning"

| **Date** | **Animal** | **Age** | **Current (uA)** | **Duration (min)** | **Voltage Related Notes** |
|---|---|---|---|---|---|
| 180702 | Sheep | Adult | 510 | 20 | no current delivered (faulty setup) |
| 180702 | Sheep | Adult | 480 | 20 | 90 V (etching; less than 30 s), 22 V |
| 180702 | Sheep | Adult | 250 | 10 | 90 V (etching; few seconds), 16 V |
| 180716 | Pig | Juvenile | 180 | 2 | immediate voltage drop |
| 180716 | Pig | Juvenile | 180 | 2 | immediate voltage drop |
| 180716 | Pig | Juvenile | 180 | 2 | immediate voltage drop |
| 180716 | Pig | Juvenile | 180 | 2 | immediate voltage drop |
| 180716 | Pig | Juvenile | 180 | 2 | immediate voltage drop |
| 180716 | Pig | Juvenile | 180 | 2 | immediate voltage drop |
| 180727 | Pig | Juvenile | 180 | 2.2 | $< 15$ V |
| 180727 | Pig | Juvenile | N/A | 5.5 | N/A |
| 180727 | Pig | Juvenile | 180 | 2 | 15 V average |
| 180727 | Pig | Juvenile | 180 | 2 | $< 15$ V |
| 180727 | Pig | Juvenile | N/A | 6 | N/A |
| 180727 | Pig | Juvenile | 180 | 1 | $< 15$ V |
| 180727 | Pig | Juvenile | 180 | 1 | $< 15$ V |
| 180727 | Pig | Juvenile | 180 | 1 | $< 15$ V |
| 180727 | Pig | Juvenile | 180 | 1 | $< 15$ V |
