## Supplementary File 2 for "Neuroelectrophysiology-Compatible Electrolytic Lesioning"

| Date | Animal | Sex | Age (wks) | Current (uA) | Duration (min) | Voltage Related Notes |
|---|---|---|---|---|---|---|
| 190215 | Pig | F | 13 | 150 | 1 | voltage railed |
| 190215 | Pig | F | 13 | 150 | 1 | < 15 V |
| 190215 | Pig | F | 13 | 240 | 1 | voltage railed |
| 190215 | Pig | F | 13 | 240 | 1 | < 15 V |
| 190215 | Pig | F | 13 | 240 | 2 | < 15 V |
| 190215 | Pig | F | 13 | 150 | 2 | < 15 V |
| 190215 | Pig | F | 13 | 150 | 5 | < 15 V |
| 190215 | Pig | F | 13 | 185 | 10 | < 15 V |
| 190326 | Pig | F | 10 | N/A | N/A | N/A |
| 190326 | Pig | F | 10 | 100 | 1 | < 15 V, briefly 30 V, < 15 V |
| 190326 | Pig | F | 10 | 100 | 0.5 | < 15 V |
| 190326 | Pig | F | 10 | 100 | 0.25 | voltage railed |
| 190326 | Pig | F | 10 | 100 | 1 | < 15 V |
| 190326 | Pig | F | 10 | 100 | 1 | < 15 V |
| 190326 | Pig | F | 10 | 50 | 0.5 | < 15 V |
| 190326 | Pig | F | 10 | 50 | 1 | < 15 V |
| 190326 | Pig | F | 10 | 150 | 0.5 | < 15 V |
| 190326 | Pig | F | 10 | N/A | N/A | N/A |
| 190416 | Pig | F | 16 | 50 | 0.5 | < 15 V |
| 190416 | Pig | F | 16 | 50 | 1 | < 15 V |
| 190416 | Pig | F | 16 | N/A | N/A | N/A |
| 190416 | Pig | F | 16 | 100 | 0.5 | < 15 V |
| 190416 | Pig | F | 16 | 100 | 1 | voltage railed |
| 190416 | Pig | F | 16 | 50 | 0.5 | < 15 V |
| 190416 | Pig | F | 16 | N/A | N/A | N/A |
| 190416 | Pig | F | 16 | 50 | 1 | < 15 V |
| 190416 | Pig | F | 16 | 100 | 0.5 | < 15 V |
| 190416 | Pig | F | 16 | 100 | 1 | < 15 V |
| 190513 | Pig | F | 10 | 50 | 1 | < 15 V |
| 190513 | Pig | F | 10 | 50 | 1 | < 15 V |
| 190513 | Pig | F | 10 | N/A | N/A | N/A |
| 190513 | Pig | F | 10 | 150 | 2 | voltage railed |
| 190513 | Pig | F | 10 | 150 | 1 | < 15 V |
| 190513 | Pig | F | 10 | N/A | N/A | N/A |
| 200203 | Pig | F | 9 | 450 | 0.2 | |
| 200203 | Pig | F | 9 | 450 | 0.5 | |
| 200203 | Pig | F | 9 | 450 | 0.2 | |
| 200203 | Pig | F | 9 | 450 | 0.2 | |
| 200203 | Pig | F | 9 | 450 | 0.2 | |
| 200203 | Pig | F | 9 | 450 | 0.2 | |
| 200203 | Pig | F | 9 | 450 | 0.2 | voltage railed |
| 200203 | Pig | F | 9 | 450 | 0.2 | |
| 200203 | Pig | F | 9 | 350 | 0.2 | < 15 V |
